# Selection history alters fitness across ecological interactions in a fly–parasitoid system

**DOI:** 10.64898/2026.09.20.752945

**Authors:** Vrinda Ravi Kumar, Juan Esteves, Vincent Montbel, Somayeh Rasouli-Dogaheh, Varvara Fedorchenko, Jan Hrček

## Abstract

Species interactions shape ecological dynamics, but adaptation to one interaction can trade off with others. Experiments that measure such evolved fitness changes across multiple interactions remain rare. We used experimental evolution to test whether recent selection under one ecological interaction changes fitness under other interactions in a fly–parasitoid system. We evolved a genetically diverse *Drosophila melanogaster* population for 15–17 generations under three regimes: benign conditions (control), high intraspecific competition, and infection by the parasitoid wasp *Leptopilina boulardi*. We then measured fly fitness (egg-to-adult survival and subsequent reproduction) under benign conditions, high intraspecific competition, and infection by three other parasitoid species and an additional conspecific *L. boulardi* strain. Selection history strongly influenced fitness across interactions. Populations maintained under high larval density showed reduced survival under infection by *L. boulardi* and *L. victoriae*, indicating a trade-off between past selection for competition and defence against some parasitoids. Populations evolved under parasitoid infection by *L. boulardi* showed increased resistance to a conspecific strain but reduced fitness when infected by another species, *L. heterotoma*. Our results show that rapid evolution in response to a single interaction can reshape multiple ecological interactions through fitness trade-offs, highlighting how recent evolutionary history may influence community dynamics and invasion susceptibility.

## Introduction

Trade-offs between species interactions (such as competition, predation and parasitism) are considered key drivers of species coexistence and community dynamics [1–3]. The species interactions can often change systematically across seasons, with a major impact on community composition and persistence [4,5]. Changes in interactions may also happen much more abruptly, for instance, when ecosystems cross tipping points [6,7], or during species extinctions or invasions [8–10]. Such abrupt transitions may drive the rapid evolution of interaction-related traits which can in turn feed back to community dynamics. Theory explores the community-level impact of evolutionary trade-offs in interaction traits [11–14]. Understanding such trade-offs empirically is an important next step to predict community composition and stability. It is even more critical in the context of increasingly unpredictable climate variations and anthropogenic activity in rapidly urbanizing areas, which may disrupt natural species interactions and lead to complex community restructuring [15–18].

Despite theoretical work, several empirical gaps remain in how interactions within a community trade off [19,20]. The competition-defence trade-off is commonly studied empirically. This trade-off is typically between competitive ability and defence against predators, pathogens or parasites, although it is not always reciprocal and symmetric (examples: bacteria-protozoa [21,22]; bacteria-phage [23]; algae-rotifer [24]; fruit fly-parasitoid wasp [25,26]; fish-predator [27]; plant-predator [28]). We argue here for two critical gaps limiting the extension of these measurements to broader community contexts. First, overall fitness (rather than the specific trait value of any competition or defence-related phenotype) will directly determine species abundance and therefore, any downstream community consequences, as has been argued before [29]. Since fitness integrates the effect of multiple phenotypes, and adaptation is often polygenic [29], it is critical to quantify the impact of trade-offs on overall fitness under a given interaction [30]. This is typically already done in studies on microbial or near-microscopic species [21–24], but rarely in macroscopic organisms.

Second, evolutionary trade-offs between species interactions for macroscopic organisms have usually been studied using artificial selection for a specific trait [31]. Artificial selection is experimentally imposed by measuring a specific trait of interest and only propagating individuals above a chosen threshold of the trait value, similar to selective breeding in domesticated organisms [32]. There are clear advantages to using such artificial selection for estimating quantitative genetic parameters, generating strong predictable responses to selection, and having direct control over selection intensity. In contrast, in experimental evolution, populations are exposed to controlled environmental conditions and which individuals survive and reproduce is a direct function of their overall fitness in those environments. Experimental evolution is thus better suited to study adaptation in ecological contexts because selection arises from the environmental conditions themselves rather than being artificially imposed by the experimenter [31,33]. Further, the method has clear consequences for the detection of trade-offs and their effect size. For instance, one study reported that experimental evolution resulted in about half the increase in parasitoid resistance compared to artificial selection in another comparable study [34]. Finally, while many studies characterise this trade-off within a single species interaction (one competitor and one predator, pathogen or parasite; but see [26] for a total of six studied species), it is critical to extend the number of studied interactions to extrapolate to a multispecies context. In sum, there is a critical convergence of methodological gaps in our empirical understanding of how rapid evolution reshapes ecological interactions in a community context.

Fly-parasitoid systems are a great model to study interaction-related trade-offs and their community-level impacts because parasitoids often impose strong selection on flies, which, in turn, can rapidly evolve defences [25,26,35,36]. Parasitoids employ diverse infection strategies [37], potentially eliciting different resistance mechanisms [38]. Simultaneously, flies often face intense competition during the larval stage, and competitive ability can evolve [39]. Previous experiments have documented that when parasitoid resistance is artificially selected for, competitive ability reduces in laboratory populations [25,40], broadly suggesting genetic constraints on simultaneous optimization of both traits. However, one study which selected for increased competitive ability in *Drosophila melanogaster* found that parasitoid resistance increased, highlighting that the detection and direction of this trade-off is highly context-dependent [41], further underscoring the need for using ecologically-relevant selection experiments and measurements that can be extrapolated to the community context. Despite the relevant background studies highlighted above, whether fitness under different parasitoid and competitive interactions trade off with each other remains untested, to the best of our knowledge, as does the generality of such effects across other parasitoid species.

Here we test whether population-level fitness components (survival and subsequent reproductive ability) in the presence of parasitoids trade off with overall fitness under intraspecific competition. We used populations from a recent experimental evolution study using a genetically diverse *D. melanogaster* population subjected to three different selective regimes (benign control conditions, high larval competition, and exposure to the parasitoid wasp *Leptopilina boulardi*) for 15-17 generations in five replicates per selection regime. Our experimental evolution approach allowed natural selection to operate across multiple fitness components simultaneously. We tested evolved populations for overall fitness (survival and reproduction) under crowded larval conditions and against three other parasitoid species ranging in relatedness to the focal *L. boulardi,* as well as a different conspecific *L. boulardi* strain. This design allows us to quantify the emergence and generality of evolutionary trade-offs across fitness under multiple ecological interactions simultaneously.

## Methods

### Ancestral fly stock

We initiated the selected populations from a genetically variable starting population of *Drosophila melanogaster* created by mixing 90 isofemale lines obtained from the DrosEU consortium (Figure 1), to sample ecologically relevant levels of variation. The lines were collected from nine locations in Europe during the summer and fall of 2018 [42]. To create the starting population, we mixed five males and five females from each line in a cage (46 x 30 x 23cm) with mesh-covered cutouts for ventilation. This mixed population recombined and stabilized in the laboratory for 35 generations prior to the initiation of selection populations, following a 14-day discrete generation cycle. Each generation was initiated with eggs collected between 3–5 days post-eclosion. Eggs were dispensed at a fixed volume of 21uL (protocol described in [43]) per bottle containing 40 mL of cornmeal-yeast-sugar food. This egg volume averages ∼400 fly eggs, leading to a stock larval density of 10 eggs/mL of food. The ancestral stock and all derived selection populations were maintained at 25 ± 1°C with 60% RH at a 12 hour light/dark photo-period cycle.

**Figure 1.**
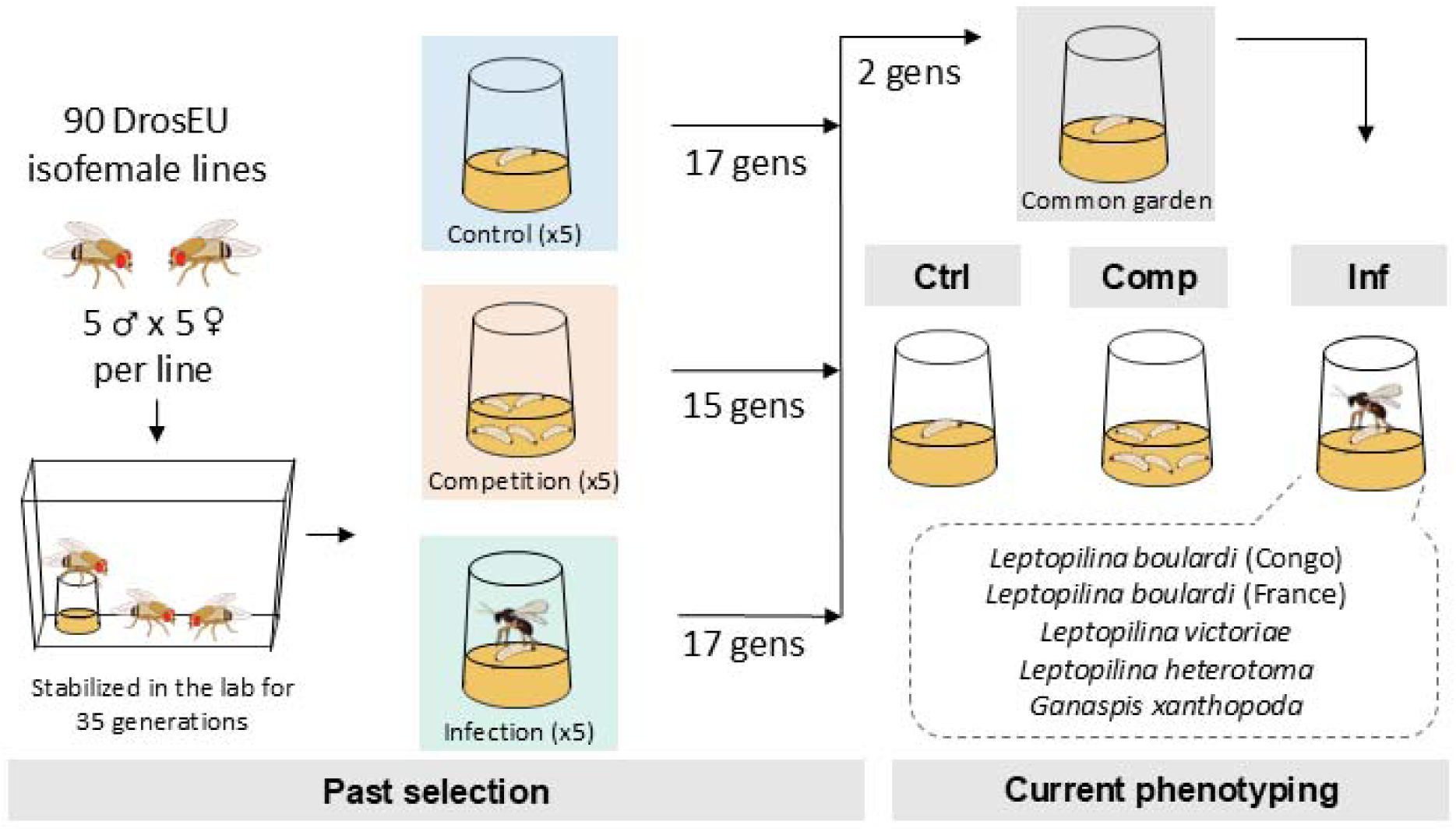
Schematic of the past selection regimes and the current phenotyping experiment. During phenotyping, Ctrl = low larval density, uninfected; Comp (fitness under competition) = high larval density; Inf (fitness under parasitoid infection) = infected at low larval density.

### Parasitoid wasp origin and maintenance

Parasitoid wasps used in the phenotyping assays were maintained on another outbred population of *D. melanogaster* under the same environmental conditions. We used a total of four species of larval parasitoid wasps from the family Figitidae, which are koinobionts (their hosts continue to feed and develop after parasitisation). The focal strain used for both past selection and current phenotyping was *Leptopilina boulardi* collected in Brazzaville, Republic of Congo [44]. The other strains varied in relatedness to the focal strain and were used for the current phenotyping only: another strain of *Leptopilina boulardi* from Lyon, France [45], *Leptopilina heterotoma* from Trentino, Italy (by Miguel González Ximénez de Embún), *Leptopilina victoriae* from Hawaii [46], and *Ganaspis xanthopoda* from Uganda [46].

### Selection history

After allowing for recombination in the ancestral fly stock, we created derived, replicated selection populations under three regimes - 1) “Control” identical to the ancestral stock regime, 2) “Competition” with five times the control larval density during development resulting in high larval competition, and 3) “Infection” by the Congo strain of *Leptopilina boulardi* where 48-hour old fly larvae were infected at a fixed infection strength of 1 female parasitoid per 100 eggs (Figure 1). Selection treatments were maintained at large population sizes (over 1500 individuals) during selection to minimize stochasticity in the selected phenotypes. We used experimental evolution rather than artificial selection, which allows for a broader range of evolved responses (as opposed to experimenter-scored phenotypes). These selection populations were part of a larger laboratory evolution experiment (Montbel *et al*., in prep), with relaxed and test individuals generated separately for this experiment. All selection populations were initiated simultaneously with six replicates per selective regime. For this experiment, we included five replicate populations per regime with the most consistent survival in earlier generations.

The populations were maintained on a discrete generation cycle, set to three days after peak adult emergence time under each of these regimes - 14 days for Control and Infection, and 15 days for Competition. Prior work shows that flies reach peak fecundity at this time [47]. To start the next generation, we allowed overnight egg-laying on a yeasted 4% agar plate to stimulate egg laying.

The next day, we dispensed the eggs collected from these plates into fresh food bottles using the same method as the ancestral stock described above. The egg volume in Control and Infection regimes was identical to the stock regime (21 uL eggs in 40 mL food) and Competition was dispensed at 5 times the egg volume (105 uL eggs in 40 mL food). Infection treatment was subsequently infected 48 hours after dispensation with 4 fertilized female wasps (between 3-7 days post eclosion) per bottle. Every generation, for each replicate, we generated 8 bottles for Control and Infection, and 2 bottles for Competition. We aimed for a minimum adult fly density of 1500 individuals to minimize drift under the selective regimes.

### Phenotyping experiment

After propagating the populations for 17 generations for “Control” and “Infection” and 15 generations for “Competition” under their respective selection regime, five replicate populations per regime were relaxed for two generations under benign common garden conditions to minimize the impact of phenotypic plasticity. We collected third generation relaxed eggs and manually counted them in PBS for the phenotypic assays, creating six replicate vials per phenotyping treatment. All populations were phenotyped in vials containing 6 mL of fly food. The phenotyping treatments intentionally closely resemble the past selection regimes described above. We therefore refer to the current phenotyping treatments with abbreviations to clearly distinguish them from the past selection regimes. We established a total of seven phenotyping treatments - “Ctrl”, “Comp” and “Inf” (with five sub-treatments within “Inf” for each of the tested wasp strains; Ctrl - 60 eggs; Comp - 300 eggs; Inf - 60 eggs infected by female wasps after 48 hours; Figure 1). The phenotyping experiment consisted of a total of 59400 manually counted eggs (300 eggs/vial for Comp, and 60 eggs/vial for the other 6 treatments, across 630 phenotyping vials (6 replicate vials per treatment-population combination in 7 treatments for each of 15 replicate selection populations)).

We used different infection strengths for the five wasp strains, calibrated to maximise the probability of picking up any evolved changes in resistance based on a series of pilot experiments. For the strains with lower virulence (*L. boulardi* (Congo), *L. heterotoma*, and *L. victoriae*), we exposed host larvae to two fertilized female wasps per 60 eggs, while for strains with high virulence (*L. boulardi* (France) and *Ganaspis xanthopoda*) we exposed host larvae to one fertilized female wasp per 60 eggs.

Viability was measured as egg-to-adult survival and was quantified as the proportion of adult flies that emerged from the eggs put in the vial. Fly emergence was scored 3-4 days after complete eclosion of adult flies, 12-15 days after egg collection. Immediately after measuring viability, we transferred 3 random females per viability vial to a new vial to measure fertility. These females were removed from the vial 72 hours later. Fertility was measured as the number of adult offspring flies per vial, and the counts were performed two weeks after the transfer. The viability and fertility phenotypic measures are intentionally broad to integrate the net effect of multiple adaptations to the selection regime into two overarching fitness measures. Further, this method allows us to directly compare effects across different interspecific interactions like intraspecific competition and parasitoid infection.

### Statistical analysis

To account for baseline differences between the evolving selection populations, we normalized all fitness measurements by dividing them by the mean fitness value of the same population under control conditions (viability and fertility). This approach allowed direct comparison of stochastically diverged populations with different baseline fitness levels under each experimental condition. We then used these normalized viability and fertility values from the core treatments only (Ctrl, Comp, Inf) to first test whether the populations responded to the past selective regimes. Then we compared across the entire dataset, also including the four other parasitoid populations. We tested for differences using Gaussian linear mixed-effects models (LMMs) in the lme4 package in R (version 4.4.1). We modelled both fitness variables as a function of selection regime, phenotyping treatment and their interaction, with population replicate as a random effect (formula: normalized fitness metric ∼ selection regime × treatment + (1|population replicate)). We fit these LMMs to the entire dataset and visualize replicate-level differences in Figure 2 for Comp and Inf (the conditions under selection). While the mixed model fit predicted 80.3% and 44.6% of the variance in the whole dataset, fixed effects alone explained 79.4% and 33.9% for normalized viability and fertility, respectively. Finally, model comparison using AIC and BIC indicated no improvement over a fixed-effects model. Thus, we only report statistical results from the fixed-effects linear model in the manuscript, effectively combining all the population replicates within the selection regime. From these models, we extracted estimated marginal means using the emmeans package and performed pairwise comparisons of relative (normalized) viability and fertility between selection regimes, within each treatment. All pairwise comparisons within the full model were adjusted for multiple testing using Tukey’s HSD.

**Figure 2.**
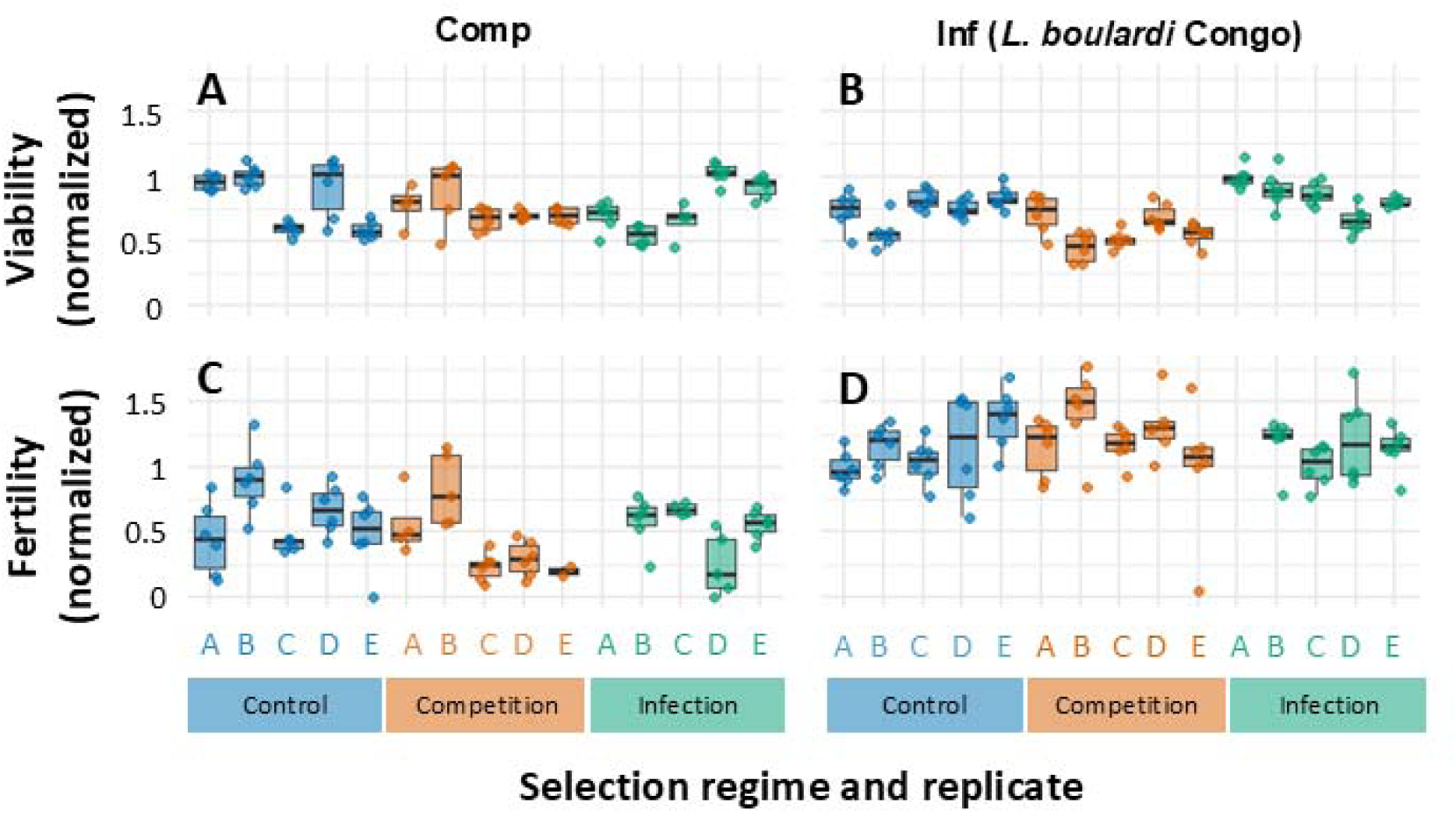
(A, B) Normalized viability and (C, D) fertility for all five replicate selected populations (replicates A-E, blue = Control; orange = Competition; green = Infection) measuring 2 traits – (A, C) fitness under competition (“comp”), (B, D) fitness under parasitoid infection with *Leptopilina boulardi* (Congo strain) which was also used to impose selection in the Infection selection regime. We visualize all the replicate populations in this figure separately to represent inter-population variation, but all statistics in the main text and SI report trends across selection regimes (all replicates combined).

## Results

### Evolved fitness changes and trade-offs in response to selection history

We first explored evolutionary responses in the fitness-related phenotypes we directly selected for under high larval density (Competition) and infection by the Congo strain of *L. boulardi* (Infection) to understand the baseline impact of selection history (Figure 2). The response of replicates to selection treatments was largely consistent, as indicated by lack of improvement of the model fit by including replicate as a random factor. Infection populations (evolved in the presence of parasitoids) showed a marginally significant 10% increase in relative (normalized) viability in response to infection after 17 generations (Table S1, linear model, p = 0.055; relative to control). The Competition populations (evolved under high larval density) did not evolve higher viability under high larval density in our experiment (Table S1, linear model, p = 0.125). Unlike viability, fertility did not respond to the selection for the populations phenotyped under their respective regime conditions (Table S2, linear model, p > 0.05 for both).

Interestingly, we found evidence for the trade-off between infection by *L. boulardi* (Congo) and selected high larval density (Table S1, linear model, Figure 3A). Competition populations showed a strong 16% drop in relative viability under infection by *L. boulardi*, indicating that their parasitoid resistance had dropped as a correlated response to selection under high larval density (Table S1, linear model, p = 0.003). However, we did not find evidence for the trade-off in the reverse direction - Infection populations did not show altered fitness relative to control under high larval competition (Table S1, linear model, p = 0.425). We also did not find any changes in fertility across selection regime - treatment combinations (Table S2).

**Figure 3.**
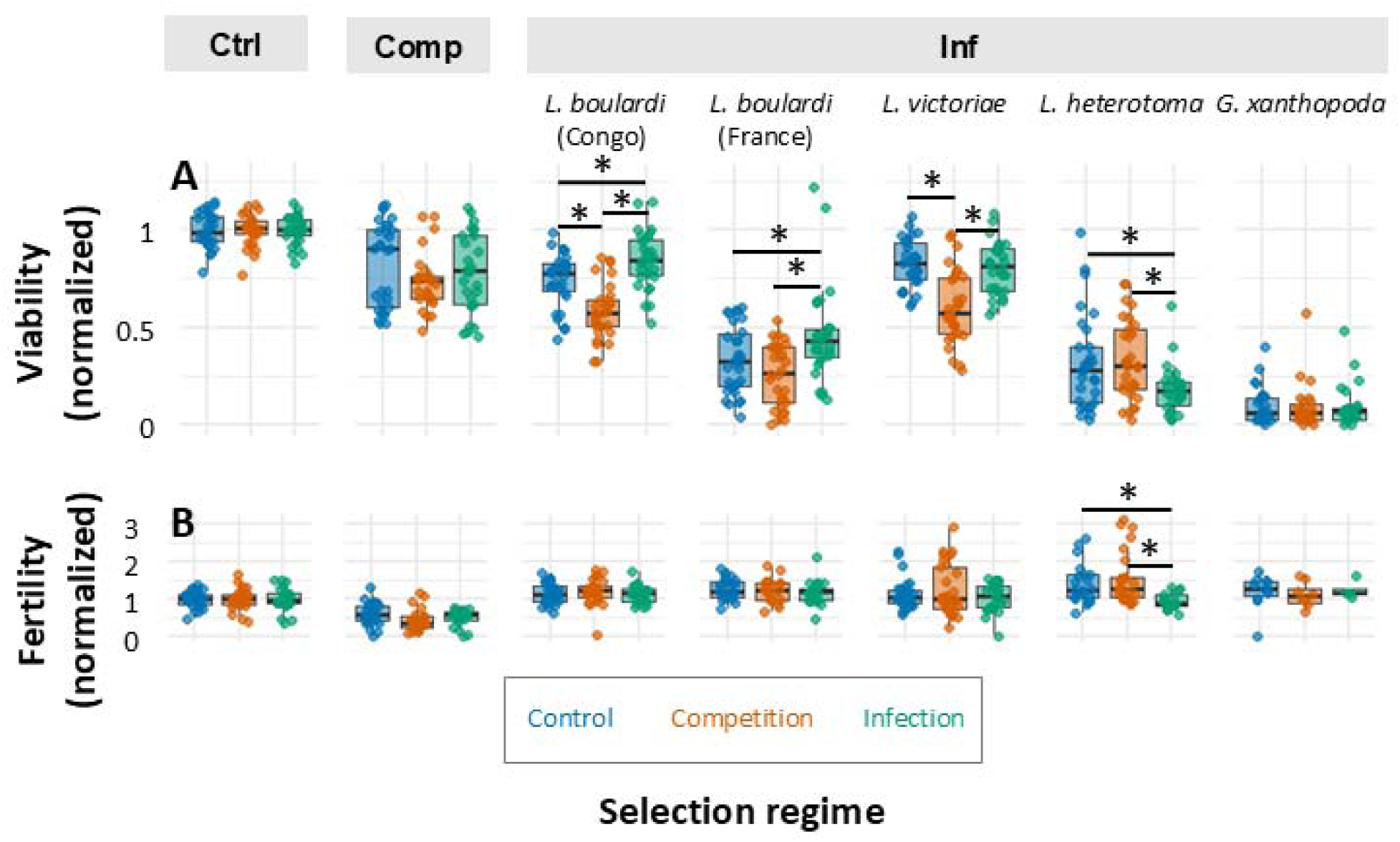
Normalized (A) viability and (B) fertility boxplots for the experimental phenotypes – Ctrl (low density), Comp and Inf by *Leptopilina boulardi* (Congo strain), *L. boulardi* (France strain), *L. victoriae*, *L. heterotoma*, and *Ganaspis xanthopoda* (Blue = Control; Orange = Competition; Green = Infection). Data are combined across five replicates for each selection regime. Asterisks indicate statistically significant comparisons (p < 0.05) for estimated marginal means from a linear model for normalized viability and fertility (full statistics reported in Table S3-S4). P-values have been corrected for multiple comparisons within a panel using Tukey’s HSD.

Our experimental conditions were initially calibrated to be stressful for the ancestral population in a series of pilot experiments. We tested whether they remained challenging to the evolved flies as a loss of experimental stress would reduce our ability to detect trade-offs. We found significant evidence that the experimental conditions remained challenging to the test flies. Being exposed to high larval density reduced relative fly viability by 18% (Table S1, linear model, p < 0.001) across all selected populations. Infection with *L. boulardi* Congo also resulted in a strong 26% decrease in relative viability (Table S1, linear model, p < 0.001). Relative fertility strongly dropped by 40% across selection regimes when the females developed under high larval density (Table S2, linear model, p < 0.001), and increased by 14% when infected by parasitoids (Table S2, linear model, p = 0.044).

### Competition - defence trade-off in other parasitoids

We reported above that populations evolved under high larval density showed reduced resistance to a Congo strain of *L. boulardi* (Figure 3A, Table S1). To test whether evolving in response to crowded conditions might leave fly populations vulnerable to other parasitoids *D. melanogaster* may encounter, we tested all Competition populations against a panel of four parasitoids varying in relatedness to our ‘focal’ parasitoid strain of *L. boulardi* (Congo). We first tested comparisons between Control and Competition populations from the full model to assess whether developing in high larval density imposes an additional cost on being infected by parasitoids. We found that this trade-off did not extend to a different strain of *L. boulardi* from France (Table S3, linear model, p = 0.184), the closest relative of *L. boulardi* Congo. However, Competition populations showed a strong 22% reduction in relative viability on infection by one out of the three newly tested parasitoid wasp species - *Leptopilina victoriae* (Figure 3A, Table S3, linear model, p < 0.05), but not against the other species (Table S3, p > 0.05 for all others).

We also tested this trade-off using a more extreme comparison - Competition populations vs the Infection populations. Populations across these selective regimes are likely more diverged from each other along this trade-off than Control vs Competition regimes in the same short experimental timeframe, increasing our detection power. As expected, we found strong effects, further supporting the existence of this trade-off. Competition populations showed significantly higher mortality than Infection populations on infection with *L. boulardi* Congo (by 26% relative viability), but also with *L. boulardi* France (by 20% relative viability), and *L. victoriae* (also by 20% relative viability) (Figure 3A, linear model, p < 0.05 for all, Table S3). On the contrary, Competition populations had higher survival and fertility than Infection populations on infection by *L. heterotoma* (by 16% relative viability and 60% relative fertility) (Table S3-S4).

### Cross-resistance to other parasitoids

Finally, we tested whether populations evolved in the presence of one parasitoid species (Infection populations) were more or less vulnerable to new parasitoids relative to Control populations. We found divergent patterns, likely indicating the complexity and diversity of resistance mechanisms. Populations selected for resistance to *L. boulardi* (Congo) also showed increased cross-resistance against its conspecific strain *L. boulardi* (France) by a 12% increase in relative viability (Figure 3A, Table S3, linear model, p < 0.05). We did not find transfer of resistance to *L. victoriae* and *G. xanthopoda*. On the contrary, evolved Infection populations showed strongly reduced fitness (14% in viability and 45% in fertility) on infection with *L. heterotoma*, relative to Control populations (Figure 3A-B, Table S3-S4). This implies a trade-off between evolved resistance to *L. boulardi* and fitness when attacked by *L. heterotoma*.

## Discussion

We empirically tested whether recent selection history (15-17 generations, ∼7-8 months) on a focal species can alter other species interactions in a community. Our study provides a new test in an insect community consisting of the fruit fly *D. melanogaster* and four parasitoid wasp species. We directly measure fly survival and reproduction within a given interaction (as discussed in [30]), rather than a specific phenotype that may only partially contribute to either of these. Our experiment uncovered strong and direct downstream fitness effects of past selection (on both fitness under high larval density and on infection by *L. boulardi*) across multiple interspecific interactions. Past selection under high fly larval density strongly impacted fly fitness after infection with parasitoid wasps for three out of four tested wasp species, even though it had no measurable effect on fitness under high density itself in our experiment. Parasitoid resistance evolved within a few months, even with experimental evolution instead of artificial selection. Prior work has demonstrated that experimental evolution can result in about half the increase in resistance compared to artificial selection [25,34]. Selection under wasp infection with *L. boulardi* (Congo) altered fly interactions with wasps for three out of five tested wasp species or strains. These effects were measured after two generations under common garden conditions, indicating robust evolutionary changes in fitness that may influence population dynamics and subsequently, longer-term community dynamics. These findings are direct empirical evidence that past selection on one species interaction can reshape multiple interspecific interactions in a community via fitness trade-offs.

### Trade-off between intraspecific competitive ability and parasitoid resistance

One of the main trade-offs between species interactions that we tested was between host intraspecific competitive ability and parasitoid resistance. Past selection under high larval density reduced *D. melanogaster*’s survival under infection with two parasitoid species - *L. boulardi* (Congo) and *L. victoriae*. The converse was not true in our experiment – past selection under parasitoid infection did not significantly change high-density survival. However, Competition and Infection populations also diverged significantly in fitness under infection by two additional parasitoid strains/species, which indicates the existence of an underlying trade-off. The strength of these fitness differences depends on the parasitoid - strong in some (as much as a 26% change in viability or a 60% change in fertility), and absent (or not yet detectable) in others. We conclude that the trade-off between survival under high competition and parasitoid attack can partially appear at least as early as after 17 generations of evolution. Our findings are consistent with the existence of a broader symmetric trade-off between intraspecific competitive ability and parasitoid resistance in *D. melanogaster*. Evolution along this trade-off axis may generate different levels of invasion susceptibility by a new parasitoid species in an established insect community with *D. melanogaster*.

This trade-off in fruit flies (also in other *Drosophila* species) has been partially established by other studies, most of them using artificially selected flies. Our study is novel in this regard - it tests whether Competition and Infection populations derived from the same initial population show fitness divergence. Influential work found that artificial selection for encapsulation ability against *Asobara tabida* and *L. boulardi*, a cellular immune response that surrounds and melanizes parasitoid eggs, reduced competitive ability in *D. melanogaster* [25,26,35,40], but also across two other *Drosophila* species [26]. McGonigle *et al*. [26] also found that evolved parasitoid resistance to *L. boulardi* was rapidly lost after artificial selection for encapsulation was switched to experimental evolution under high density, possibly due to this trade-off. Results from the only two experimental evolution studies on fruit flies that we are aware of are mixed. A study that experimentally evolved *D. melanogaster* under high larval density found increased (rather than decreased) correlated parasitoid resistance to *A. tabida*. The authors note the complexity of selecting for competitive ability, and caution their results may be due to improved wound healing [41]. Notably, they also did not find increased competitive ability in their evolved populations when using the same assay we did (but did find higher competitive ability using a different assay). Competitive ability measurements are highly sensitive to assay choice [48,49]. This likely explains why we did not detect evolved changes in competitive fitness in our Competition populations, since phenotypic data under parasitoid infection suggests that Competition populations did diverge from the Control. Another study found that evolved resistance to *A. tabida* resulted in lower feeding rate [34] - a trait associated with fly populations maintained at low density [50] - widely recognized as a strong correlate of larval competitive ability in *Drosophila*. In non-*Drosophila* insects too, evolved resistance to a virus in the Indian meal moth trades off with host development time, which will likely impact growth rates [51].

Zooming out, trade-offs between competitive ability under resource limitation and defence against natural enemies are phylogenetically widespread beyond insects. In unicellular algae, experimentally evolved strains of *Chlorella vulgaris* and *Chlamydomonas reinhardtii* also diverge along the competition-defence axis [2,52]. In a plant pathogen, phage-driven resistance trades off with resistance to a competitor bacteria [53]. In plants, a trade-off between constitutive resistance to herbivory and competitive ability has been demonstrated across 58 species from 15 families [54]. Interestingly, the trade-off is driven by wild (and not domesticated) plant species, indicating it may be maintained by evolutionary forces rather than a fixed physiological constraint.

### Cross-resistance across parasitoids

We further directly tested whether evolved parasitoid defences in the host were generalizable across a breadth of natural enemies. We found a range of outcomes, suggesting evolved defences are unlikely to be generalizable. Infection populations selected under *L. boulardi* (Congo) showed 12% increased viability (relative to Control populations) when infected by *L. boulardi* (France, a conspecific strain), which is comparable to the 10% viability increase observed with *L. boulardi* (Congo). However, fitness decreased under infection by *L. heterotoma*, and did not change for the two other parasitoid species (*L. victoriae* and *G. xanthopoda*). The decline in fitness under*L. heterotoma* infection is particularly noteworthy - while the 14% drop in viability is modest, the 45% fertility drop is very strong, potentially indicating a physiological trade-off between immunity to *L. boulardi* versus *L. heterotoma*. Our findings suggest that the immunological architecture underlying parasitoid resistance in *D. melanogaster* is at least partially specific. Selection under one parasitoid species can confer resistance to conspecific strains while simultaneously imposing fitness costs under attack by a different parasitoid species. If host populations evolve under locally dominant parasitoid species (e.g., *L. boulardi* in our case), they may simultaneously become more vulnerable to secondary invasion by novel or co-occurring parasitoids with different infection mechanisms.

Other studies further highlight that cross-resistance is highly context specific. Two artificial selection studies that selected for encapsulation ability in response to *L. boulardi* found large increases in resistance to *L. heterotoma* [26,35], contrasting with the fitness decrease we observed. The incongruence between our study and these studies may reflect different adaptive trajectories taken by lab populations responding to artificial selection vs experimental evolution. Experimental evolution may uncover trade-offs between different resistance strategies by exposing hidden costs that remain masked under artificial selection. Artificial selection studies on parasitoid resistance have historically used encapsulation as a readout, effective and convenient due to the ease of experimenter scoring. However, cellular immunity is only a single arm of parasitoid defence [55]. Other strategies include behavioural evasion [55] and humoral immunity [56], and a more natural selection for parasitoid resistance may involve a different balance across these immunity types. Artificial selection for encapsulation alone may drive the evolution of broadly upregulated cellular immunity that confers cross-resistance across endoparasitoid infection strategies. On the other hand, our experimental evolution protocol which exposed populations to parasitoids under more natural conditions, may have favoured a narrower, more targeted response to *L. boulardi*’s specific virulence mechanism. The strong fertility cost we observed under *L. heterotoma* infection may reflect the sensitivity of reproductive investment to this narrow targeted response to *L. boulardi*’s infection mechanism under a different immune challenge, a hypothesis that would need direct testing. Recent evidence also suggests that encapsulation and melanization may not be general defence responses against parasitoids, even within *Drosophila* [57]. Regardless of the mechanism, our results demonstrate that adaptation to one parasitoid species does not necessarily improve and can even compromise defence against another parasitoid species with a different infection strategy. This high specificity of defence against natural enemies spread across enemy guilds extends beyond *Drosophila*. For instance, in pea aphids, resistance to two parasitoid species was positively correlated, but uncorrelated between the parasitoids and a fungal pathogen [58]. In sum, host populations exposed to one natural enemy may remain entirely naive to others.

### Conclusions

Interspecific interactions, long recognized as a central focus in classical community ecology, remain underexplored in the context of rapid evolution [19,59–61]. In this context, our study strongly indicates that rapid evolution in one species has the potential to impact other members of the community via trade-offs in fitness across a range of species interactions. Since we investigated single selection pressures while populations in nature regularly encounter combinations of multiple selection pressures, our results likely represent the upper bound of interaction-related fitness evolution due to trade-offs [62]. While there is still a substantial gap between the microcosm experiment we report here and highly complex natural community dynamics, we offer some broader conclusions that may be drawn from our findings. Populations’ resilience to changes in species interactions (e.g., invasion) may be strongly influenced by their recent interactions with other species via evolution. Interactions with diverse parasitoids, in particular, might be strong drivers of trait variation in a natural community context. Despite the varied mechanisms and traits involved, our work adds to prior work broadly supporting the view that the competition–defence trade-off is a general evolutionary constraint. The ecological consequences of this evolutionary constraint spill over to two levels. Within a population, the trade-off maintains genetic polymorphism in competitive ability or growth on one side and defence on the other side [63]. Within a community, it prevents any one species from simultaneously competitively excluding other species while defending itself effectively from enemies (“being the best at everything”, [64]). Taken together, this trade-off may contribute to the maintenance of species diversity in communities.

However, neither the existence nor the direction of these trade-offs seems generalizable across species. In addition, these effects may not be direct or additive in a community context [20,65]. There are many more challenges for predicting community-level consequences of rapid evolution. We propose that (a) using experimental evolution (that allows a broader and more natural range of trade-offs than artificial selection) and (b) measuring fitness-related phenotypes with a direct impact on population growth (such as survival and fertility) are key methodological tools in this endeavour. These challenges notwithstanding, such research is particularly critical for biological control applications, invasive species management and ecosystem restoration. We found that recent evolutionary changes of a species (even within the past 20 generations) can shift species fitness under other interactions, potentially changing future adaptive trajectories and population dynamics of that species in a community context. As natural populations increasingly face unprecedented selective pressures from climate change and habitat modifications, understanding these evolutionary ripple effects through communities is key to maintaining biodiversity.

## Data availability

Data will be uploaded as Supplementary files for review and archived in Zenodo upon acceptance.

## Author contributions

Conceptualization: VRK; VM; JH

Data curation: VRK; JE

Formal analysis: VRK

Funding acquisition: JH; VRK

Investigation: VRK; JE; VM; SR; VF

Methodology: VRK; JE; VM

Project administration: VRK

Software: VRK

Supervision: JH; VRK

Validation: VRK; JE; VM; SR

Visualization: VRK

Writing – original draft: VRK

Writing – review & editing: VRK; JH; VM; JE

## Acknowledgements

We thank Andrea Weberová and Iveta Bláhová for preparing and processing the fly media, and Daria Kalendareva and Den Khvan Pak for laboratory assistance. We also thank Miguel González Ximénez de Embún, Tomáš Doležal, and Todd Schlenke for providing the parasitoid strains used in these experiments.

