## Supplementary Information for "Selection history alters fitness across ecological interactions in a fly–parasitoid system"

**Table S1.** Summary statistics from a linear model: normalized viability ~ selection regime * treatment only for the selection regimes relative to control lines. Statistically significant effects are in bold (and one marginally significant effect at p > 0.05, marked with †).

| **Factor** | **Level** | **Estimate** | **SE** | **t** | **P-value** |
| --- | --- | --- | --- | --- | --- |
| Selection regime | Competition | 0 | 0.038 | 0 | 1 |
|  | Infection | 0 | 0.0377 | 0 | 1 |
| Treatment | **Comp** | **-0.1797** | **0.038** | **-4.7258** | **<0.0001** |
|  | **Inf (*L. boulardi* Congo)** | **-0.2571** | **0.0377** | **-6.8179** | **<0.0001** |
| Interaction (Selection * Treatment) | Competition * Comp | -0.0835 | 0.0543 | -1.5381 | 0.1253 |
|  | Infection * Comp | -0.043 | 0.0538 | -0.7993 | 0.4249 |
|  | **Competition * Inf (*L. boulardi* Congo)** | **-0.1591** | **0.0533** | **-2.9843** | **0.0031** |
|  | **Infection * Inf (*L. boulardi* Congo)** | **0.1025** | **0.0531** | **1.9309** | **0.0546 †** |

**Table S2.** Summary statistics from a linear model: normalized fertility ~ selection regime * treatment only for the selection regimes relative to control lines. Statistically significant effects are in bold.

| **Factor** | **Level** | **Estimate** | **SE** | **t** | **P-value** |
| --- | --- | --- | --- | --- | --- |
| Selection regime | Competition | 0 | 0.0718 | 0 | 1 |
|  | Infection | 0 | 0.0754 | 0 | 1 |
| Treatment | **Comp** | **-0.3968** | **0.0718** | **-5.5287** | **<0.0001** |
|  | **Inf (*L. boulardi* Congo)** | **0.1441** | **0.0712** | **2.0252** | **0.044** |
| Interaction (Selection * Treatment) | Competition * Comp | -0.1711 | 0.1047 | -1.6331 | 0.1038 |
|  | Infection * Comp | -0.0873 | 0.1087 | -0.8035 | 0.4225 |
|  | Competition * Inf (*L. boulardi* Congo) | 0.0688 | 0.1006 | 0.6832 | 0.4951 |
|  | Infection * Inf (*L. boulardi* Congo) | -0.0041 | 0.1062 | -0.0384 | 0.9694 |

**Table S3.** Estimated marginal means for normalized viability across selection regime and phenotyping treatment combinations. Values represent model-adjusted means corrected for multiple comparisons. Normalized viability was modelled using a linear model: Normalized viability ~ selection regime * phenotyping treatment. Statistically significant comparisons are in bold. Df = 593.

| **Treatment** | **Contrast** | **Estimate** | **SE** | **t-ratio** | **p-value** |
| --- | --- | --- | --- | --- | --- |
| Ctrl | Control - Competition | 0 | 0.0415 | 0 | 1 |
|  | Control - Infection | 0 | 0.0412 | 0 | 1 |
|  | Competition - Infection | 0 | 0.0412 | 0 | 1 |
| Comp | Control - Competition | 0.0835 | 0.0423 | 1.973 | 0.1198 |
|  | Control - Infection | 0.043 | 0.0419 | 1.0258 | 0.5609 |
|  | Competition - Infection | -0.0405 | 0.0427 | -0.9486 | 0.6097 |
| Inf (*L. boulardi* Congo) | **Control - Competition** | **0.1591** | **0.0408** | **3.8959** | **0.0003** |
|  | **Control - Infection** | **-0.1025** | **0.0408** | **-2.51** | **0.033** |
|  | **Competition - Infection** | **-0.2617** | **0.0408** | **-6.4059** | **<0.0001** |
| Inf (*L. boulardi* France) | Control - Competition | 0.072 | 0.0408 | 1.7616 | 0.1835 |
|  | **Control - Infection** | **-0.1238** | **0.0408** | **-3.0304** | **0.0072** |
|  | **Competition - Infection** | **-0.1957** | **0.0408** | **-4.792** | **<0.0001** |
| Inf (*L. heterotoma*) | Control - Competition | -0.0154 | 0.0408 | -0.3759 | 0.9251 |
|  | **Control - Infection** | **0.1398** | **0.0408** | **3.4236** | **0.0019** |
|  | **Competition - Infection** | **0.1552** | **0.0408** | **3.7994** | **0.0005** |
| Inf (*L. victoriae*) | **Control - Competition** | **0.2229** | **0.0412** | **5.4105** | **<0.0001** |
|  | Control - Infection | 0.0241 | 0.0408 | 0.5888 | 0.8262 |
|  | **Competition - Infection** | **-0.1988** | **0.0412** | **-4.8267** | **<0.0001** |
| Inf (*G. xanthopoda*) | Control - Competition | 0.0074 | 0.0427 | 0.1726 | 0.9837 |
|  | Control - Infection | 0.005 | 0.0423 | 0.1187 | 0.9923 |
|  | Competition - Infection | -0.0023 | 0.0427 | -0.0549 | 0.9983 |

**Table S4.** Estimated marginal means for normalized fertility across selection regime and phenotyping treatment combinations. Values represent model-adjusted means corrected for multiple comparisons. Normalized fertility was modelled using a linear model: Normalized fertility ~ selection regime * phenotyping treatment. Statistically significant comparisons are in bold. Df = 453.

| **Treatment** | **Contrast** | **Estimate** | **SE** | **t-ratio** | **p-value** |
| --- | --- | --- | --- | --- | --- |
| Ctrl | Control - Competition | 0 | 0.0996 | 0 | 1 |
|  | Control - Infection | 0 | 0.1047 | 0 | 1 |
|  | Competition - Infection | 0 | 0.1047 | 0 | 1 |
| Comp | Control - Competition | 0.1711 | 0.1059 | 1.6147 | 0.2404 |
|  | Control - Infection | 0.0873 | 0.1087 | 0.8035 | 0.7011 |
|  | Competition - Infection | -0.0837 | 0.1145 | -0.731 | 0.7452 |
| Inf (*L. boulardi* Congo) | Control - Competition | -0.0688 | 0.098 | -0.7019 | 0.7625 |
|  | Control - Infection | 0.0041 | 0.1039 | 0.0392 | 0.9992 |
|  | Competition - Infection | 0.0728 | 0.1039 | 0.701 | 0.763 |
| Inf (*L. boulardi* France) | Control - Competition | 0.0287 | 0.1138 | 0.2522 | 0.9656 |
|  | Control - Infection | 0.0868 | 0.1096 | 0.7917 | 0.7083 |
|  | Competition - Infection | 0.0581 | 0.116 | 0.5007 | 0.871 |
| Inf (*L. heterotoma*) | Control - Competition | -0.1529 | 0.1109 | -1.3789 | 0.3528 |
|  | **Control - Infection** | **0.451** | **0.1247** | **3.618** | **0.001** |
|  | **Competition - Infection** | **0.6039** | **0.1215** | **4.9721** | **<0.0001** |
| Inf (*L. victoriae*) | Control - Competition | -0.1152 | 0.0996 | -1.1565 | 0.4799 |
|  | Control - Infection | 0.0998 | 0.1059 | 0.9418 | 0.6141 |
|  | Competition - Infection | 0.215 | 0.1059 | 2.0295 | 0.1063 |
| Inf (*G. xanthopoda*) | Control - Competition | 0.0903 | 0.1763 | 0.512 | 0.8655 |
|  | Control - Infection | -0.0392 | 0.2215 | -0.1769 | 0.9829 |
|  | Competition - Infection | -0.1295 | 0.2323 | -0.5572 | 0.8428 |
